# *Ex vivo* high-resolution nano-computed tomography imaging reveals spatial architecture of the adult male mouse lower urogenital tract

**DOI:** 10.1101/2025.05.25.656031

**Authors:** AR Limkar, SM Sharma, WA Ricke

**Affiliations:** Medical Scientist Training Program, University of Wisconsin-Madison School of Medicine and Publish Health, Madison, WI 53705 USA; Molecular and Cellular Pharmacology Graduate Program, University of Wisconsin-Madison School of Medicine and Public Health, Madison, WI 53705 USA; Department of Urology, University of Wisconsin-Madison, Madison, WI 53705 USA; George M. O’Brien Urology Research Center of Excellence, University of Wisconsin-Madison, Madison, WI 53705 USA

## Abstract

Benign prostatic hyperplasia (BPH) is the most common cause of lower urinary tract symptoms/dysfunction (LUTS/LUTD) in aging men. Over the past 30 years, the prevalence of BPH has increased by 122%, rising from 50.7 million cases in 1990 to 112.5 million in 2021. It is expected that this number will continue to rise over the next 15 years with the global aging population. Although mouse models are invaluable for studying human disease, gross anatomical differences between human and murine prostates complicate their translational relevance for BPH/LUTS research. The purpose of this study was to develop and validate a nanoCT-based imaging approach to enable detailed anatomical analysis of the murine lower urinary tract, with the dual goals of advancing tools for LUTD research and improving the translational relevance of mouse models to human prostate disease. Advancements in nano-computed tomography (nanoCT) have enabled high-resolution characterization of murine organ anatomy, providing new insights into how morphological differences contribute to pathology. To accomplish this, whole lower urogenital tracts from 8-week-old healthy male C57BL/6J mice and microdissected urethras were imaged on a v|tome|x M or nanotom M nano-CT system, respectively. Images were processed for 2D segmentation and subsequent 3D reconstruction using DragonFly 3D World software. Our approach enabled high-resolution visualization and characterization of the murine lower urogenital tract, including the urinary bladder, prostate lobes, seminal vesicles, and ductus deferens, as well as the microscopic ductal architecture of the prostatic urethra. The resulting 3D reconstructions preserved native anatomical relationships and allowed for comparisons between murine and human prostate anatomy. Together, these findings establish a method that can be used for assessing anatomical and morphological changes associated with LUTD development, while also highlighting key anatomical similarities that enhance the translational relevance of mouse models for human prostate disease.

## Introduction

Benign prostatic hyperplasia (BPH) arises from a loss of tissue homeostasis within the aging prostate and is the most prevalent cause of lower urinary tract dysfunction (LUTD) in older men.[1] Over the past three decades, the global burden of BPH has risen significantly, with number of prevalent cases rising from 50.7 million in 1990 to 112.5 million in 2021, an increase of 122%.[2] This trend is expected to continue alongside demographic shifts, as it is estimated that by 2050, approximately 22% of the world’s population will be over 60 years of age.[3] BPH often presents clinically as lower urinary tract symptoms (LUTS), including urinary frequency, urgency, and nocturia, which have significant impacts on quality of life and total healthcare expenditures.[4–6] It is estimated that upward of 90% of all men over the age of 80 experience some degree of prostate mediated LUTS.[7] Analysis of years lived with disability (YLD) of common urological conditions from 1990-2017 found that BPH accounts for approximately 2.4 million YLD, three times greater than that of prostate cancer.[8] As the global population continues to age, conditions like BPH/LUTS will become more clinically significant, highlighting the need for more scientific investigation. Mouse models have become indispensable for the study of human disease. In context of LUTD, sex-steroids, inflammation, mechanical insult, and natural aging have all been used to study disease development and progression in mice.[9–12] The human prostate is a single, encapsulated organ surrounded by a thick fibromuscular capsule, whereas the murine prostate comprises multiple lobes that sit freely within the peritoneal cavity.[13–15] These fundamental differences in the gross anatomy of the human and murine prostates have raised concerns about the suitability of the mouse as a model for studying prostate associated pathology.

The development of small animal, high-resolution imaging technologies have significantly advanced our understanding of anatomical similarities and differences between humans and animal models like the mouse. Techniques such as micro-computer tomograph (microCT) and magnetic resonance imaging (MRI) have been widely applied to image and reconstruct complex organs such as the murine liver, kidney, and lung. This has led to innovations in the development of therapeutics and the study of how morphological changes in pathology lead to functional decline.[16–18] Although fundamental to our understanding of the anatomical-pathological relationship, techniques like microCT and MRI are often limited by the maximum achievable spatial resolution. Alternative approaches that provide higher-resolution visualization frequently rely on 3D reconstruction of H&E-stained serial tissue sections, a process that is both time intensive and laborious.[19, 20] Nano-computed tomography (nanoCT) expands the capabilities of microCT by achieving spatial resolution down to approximately 200 nm, in contrast to the 6–12 μm range typical of microCT and the approximate 100 μm resolution limit of MRI for soft tissue visualization. NanoCT has been used to 3D reconstruct fine anatomical structures like individual pulmonary acini, cerebral vasculature, and lacunae voids within cortical bone.[21–23]

In the context of the murine lower urogenital tract, we have previously developed an MRI based method that has been used to image and reconstruct various organs including the urinary bladder, prostate lobes, and urethra. Furthermore, we have shown that this method can be used to characterize changes in urogenital tract anatomy associated with the development of LUTD.[11, 24, 25] Although faster than reconstruction of serial sections, our MRI-based method lacked the spatial resolution to appreciate the finer anatomical details such as prostatic duct counts and branching patterns. Despite these advancements, our understanding of how the unique anatomy of the mouse lower urogenital tract correlates with the human prostate remains incomplete. Here, we aim to increase our understanding of murine lower urogenital tract morphology and ultimately bridge the anatomical gap between model systems.

In this report, we describe a novel nanoCT-based approach that allows for higher-resolution visualization and reconstruction of the murine lower urogenital tract. By resolving fine anatomical structures in three dimensions, this method enables a more comprehensive assessment of murine prostate organization and its spatial relationships to the urethra and surrounding tissues. Importantly, this approach allows for precise, quantitative measurements of the urethral lumen, rhabdosphincter, and periurethral region that are directly relevant to the study of lower urinary tract dysfunction. These insights provide an important step toward improving anatomical comparisons between mouse and human prostates, ultimately enhancing the translational relevance of murine models for investigating BPH/LUTS.

## Methods and Materials

### Animals

All animal experiments were conducted under the protocols approved by the University of Wisconsin Animal Care and Use Committee (Protocol Number: M005570). All authors comply with the ARRIVE guidelines, and all methods were performed in accordance with the relevant guidelines and regulations. Male C57BL/6J mice (cat# 000664) were obtained from the Jackson Laboratory (Bar Harbor, ME). Animals were housed under standard laboratory conditions with a 12:12 light/dark cycle and provided with food and water *ad libitum*. Mice were euthanized with carbon dioxide asphyxiation followed by cardiac puncture.

### Preparation of tissues and contrast stains

Five urogenital tracts were dissected from untreated, healthy 8-week-old C57BL/6J mice as previously described.[9] Whole urogenital tracts included the bladder, seminal vesicles, prostate lobes, and urethra, with the distal ductus deferens, ureters, testes, and gonadal fat removed. Additionally, four urogenital tracts from untreated, healthy 8-week-old C57BL/6J mice were microdissected to isolate the urethra by removing the bladder, seminal vesicles, and prostate lobes.[9] Tissues were fixed in 10% neutral buffered formalin and stored in 70% ethanol until further processing. For soft tissue contrast, specimens were stained in Lugol’s iodine (10% diluted iodine-potassium iodide solution; Sigma-Aldrich, St. Louis, MO) for 24 hours at room temperature.[26] To minimize evaporation and motion artifacts during imaging, samples were embedded in 2% low-melting point agarose/PBS gel within 30 × 24 × 5 mm disposable plastic fixtures (Fisher Tissue Path, Fisher Scientific, Hampton, NH).

### NanoCT Imaging Protocol

Whole murine urogenital tracts and microdissected murine urethra specimens were immobilized inside plastic fixtures and scanned using nanoCT systems. Whole urogenital tracts were imaged using a v|tome|x M nanoCT system (Phoenix X-ray, Waygate Technologies USA, LP; Skaneateles, New York) at 100 kV and 100 µA with a 1.0 mm copper filter, an 11.39 µm voxel size, and a 500.1 ms exposure time. Three frames were averaged, and one was skipped per rotation as the sample stage rotated 360° to collect 2,220 images per scan. Microdissected urethra specimens were scanned using a nanotom M nanoCT system (phoenix x-ray, Waygate Technologies USA) at 65 kV and 250 µA with a diamond-coated tungsten target, a 0.254 mm aluminum filter, a spot size of 0, a 3.5 µm voxel size, and a 500 ms exposure time. Three frames were averaged, and one was skipped per rotation, with 2,000 images collected per scan. Image acquisition and reconstruction of raw data were performed using Datos|x 2 software (v2.8.2 for whole urogenital tracts; v2.6.1 for urethra specimens).

### 3D Reconstruction and measurements of selected anatomic features

NanoCT tomographs containing axial, coronal, and sagittal data from each scanned tissue (whole urogenital tracts and microdissected urethra specimens) were imported into DragonFly 3D World (Comet Tech, Herrengasse, Switzerland) for 3D reconstruction. Anatomical structures were identified and manually segmented using voxel thresholding based on the upper and low Otsu method. Representative 2D tomographs and 3D reconstructions are shown of a single whole urogenital tract and single microdissected urethra.

Anatomical measurements including urethral luminal area, diameter, perimeter, rhabdosphincter thickness, and prostatic duct count were taken at a standardized point within the prostatic urethra. This reference point was determined by identifying the axial slice where the rhabdosphincter fully encircled the prostatic urethra, then selecting a location 50 μm caudal to it. Urethral luminal area and perimeter were calculated using the polygon ROI tool, while luminal diameter was measured as the horizontal distance across the lumen. Rhabdosphincter thickness was determined by averaging eight measurements across its internal and external boundaries. Prostatic ducts were counted manually. Prostatic urethra length was measured in the coronal view as the distance from the start of the rhabdosphincter to the apex of the verumontanum.

## Results

Macroscopically, the murine male lower urogenital tract consists of the urinary bladder, paired seminal vesicles, urethra, and paired anterior, ventral, dorsal, and lateral prostate lobes (Fig 1). The dorsal and lateral lobes are often collectively referred to as the dorsolateral prostate.

**Fig 1.**
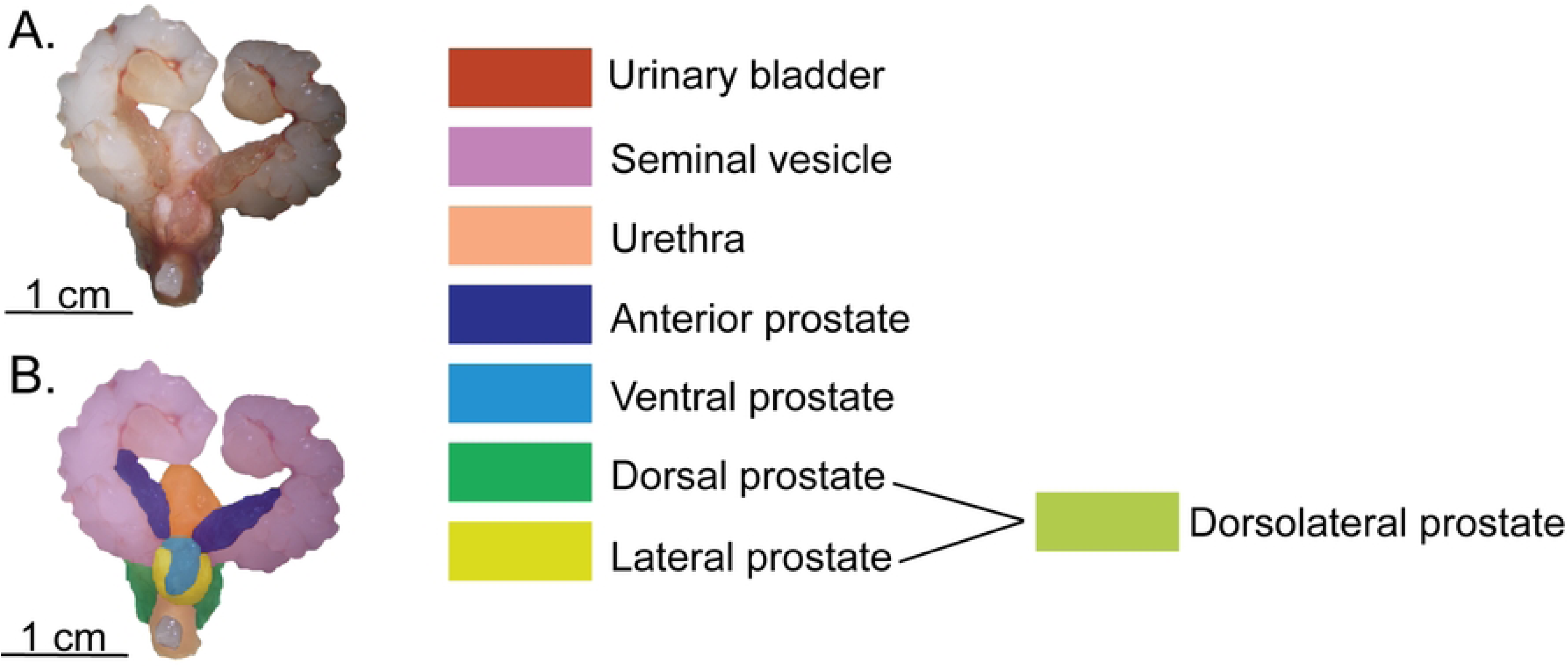
Illustration and color segmentation of the healthy 8-week-old C57BL/6J male murine lower urogenital tract. A) *Ex vivo* image of the lower urogenital tract. B) Color-coded segmentation highlighting key structures, including the seminal vesicles, urinary bladder, prostate lobes, and urethra. The color scheme will be used in all subsequent Figs.

The structural organization of the murine lower urogenital tract is illustrated by representative coronal and axial nanoCT images (Fig 2). Whole-organ scans at 11.39 μm/voxel (Figs 2A and 2B) show the relative positioning of the bladder, seminal vesicles, and prostate lobes. Manual segmentation (Figs 2C and 2D) delineates each structure segmented in the 2D plane, facilitating both quantitative and morphological analyses. Higher-resolution imaging at 3.5 μm/voxel (Fig 2E) captures fine anatomical details within the prostatic urethra, including the rhabdosphincter and periurethral prostatic ducts, with corresponding manual 2D segmentations (Fig 2F). Lugol’s iodine staining provided enhanced soft tissue contrast, allowing differentiation between glandular, muscular, and connective tissues. 2D segmentation enabled precise 3D reconstruction of each structure (Fig 2G). Together, these data establish a comprehensive anatomical framework for subsequent qualitative and quantitative analyses of individual structures in the healthy adult murine lower urogenital tract, which are detailed in the following sections.

**Fig 2.**
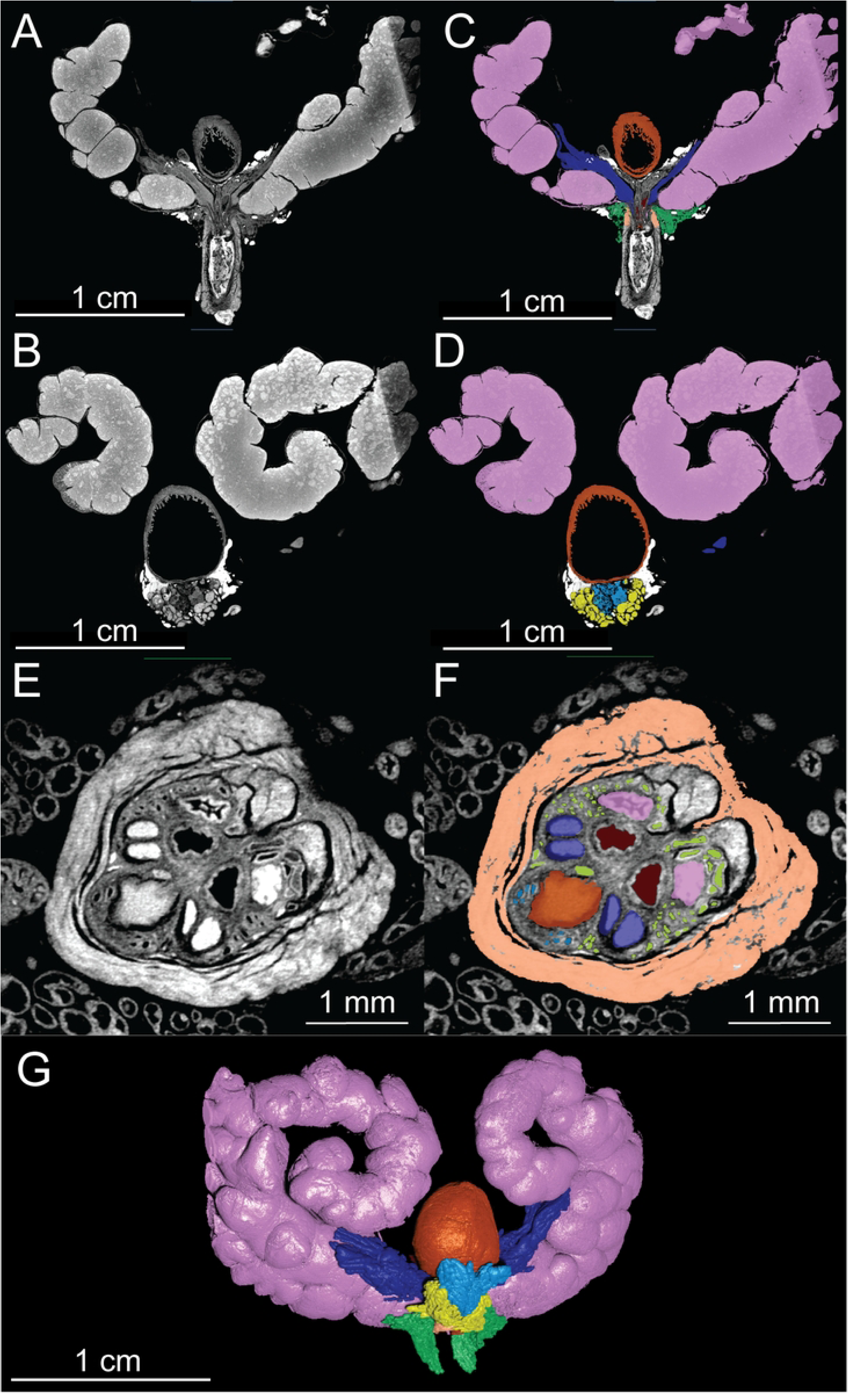
NanoCT tomographs and 2D segmentation of a healthy 8-week-old C57BL/6J male murine lower urogenital tract. A-B) Coronal nanoCT views of the intact murine urogenital tract (11.39 μm/voxel). C-D) Manual segmentation of key anatomical structures. E) Axial nanoCT image of the murine prostatic urethra (3.5 μm/voxel). F) Corresponding manual segmentation highlighting relevant structures within the prostatic urethra. G) 3D reconstruction of the intact urogenital tract. Urinary bladder (orange), seminal vesicle (pink), rhabdosphincter (light orange), anterior prostate (dark blue), ventral prostate (light blue), lateral prostate (yellow), dorsal prostate (green).

### NanoCT of the whole lower urogenital tract

NanoCT imaging at the 11.39 μm/voxel spatial resolution provided detailed visualization of the healthy 8-week-old C57Bl6/J male murine lower urogenital tract, capturing both its gross anatomical organization and fine structural features. At this lower resolution, even finer anatomy such epithelial folding patterns, large ductal architecture, and tissue boundaries were able to be visualized.

### Urinary bladder and urethral lumen

NanoCT imaging provided high-resolution visualization of the urinary bladder, enabling assessment of features relevant to lower urinary tract dysfunction (Fig 3A). Lugol’s iodine contrast effectively delineates the detrusor muscle from surrounding connective tissue and the urothelium, allowing for detailed evaluation of bladder wall architecture. Two-dimensional images reveal urothelial invaginations and areas of variable wall thickness. In the case of lower urinary tract dysfunction, acute or chronic urinary retention often results in overdistension of the urinary bladder and can lead to detrusor muscle atrophy and flattening of bladder rugae, both of which may be identified on nanoCT imaging.

**Fig 3.**
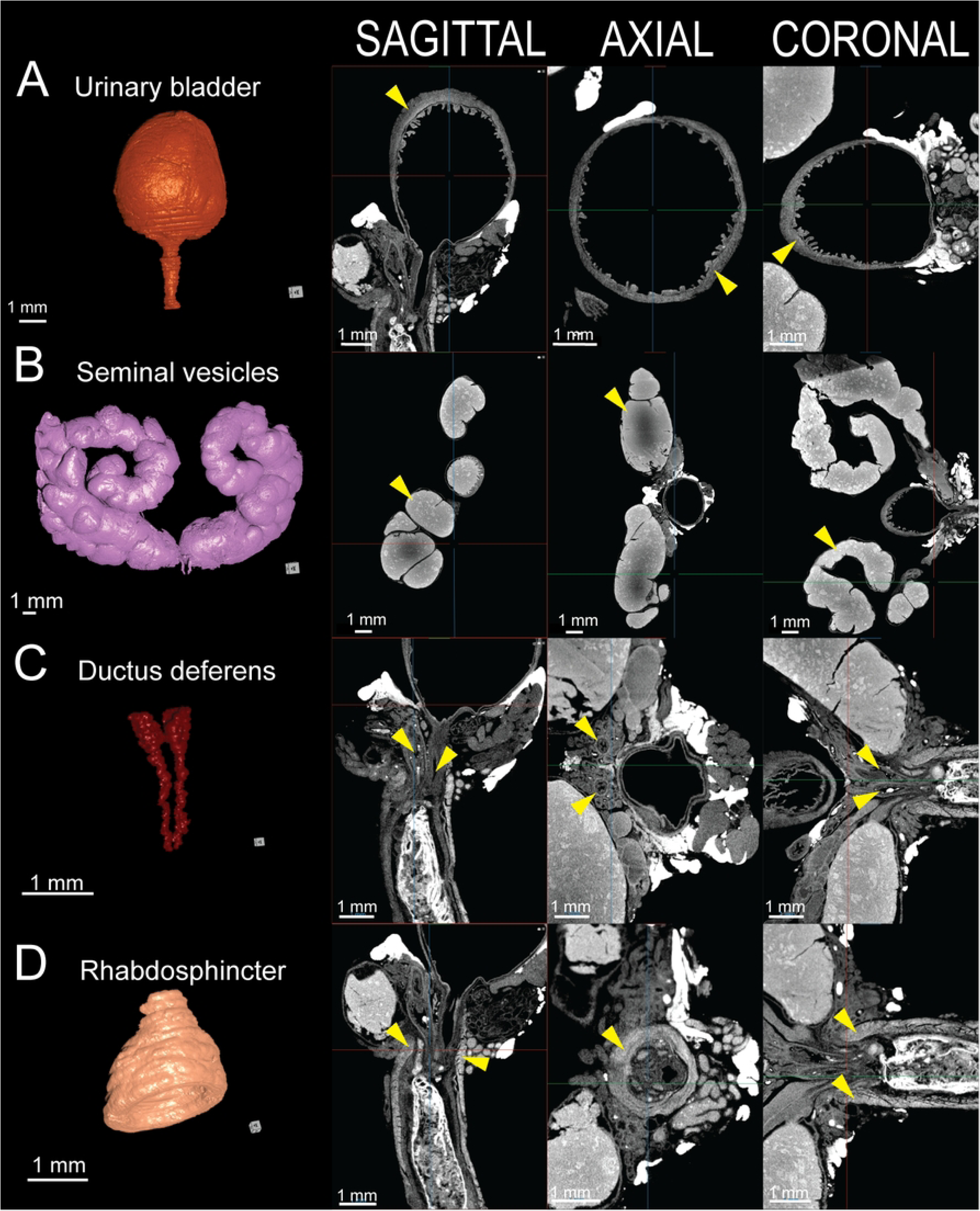
3D reconstruction and 2D tomographic views of key structures in the healthy 8-week-old C57BL/6J male murine lower urogenital tract. A) Urinary bladder. B) Seminal vesicles. C) Ductus deferens. D) Rhabdosphincter. Next to each 3D reconstruction, corresponding sagittal (left), axial (center), and coronal (right) nanoCT slices illustrate the anatomical organization. Urinary bladder (orange), seminal vesicle (pink), ductus deferens (red), rhabdosphincter (light orange).

### Seminal vesicles

NanoCT imaging enables detailed visualization of the paired seminal vesicles, revealing their complex internal architecture and luminal folds (Fig 3B). The protein-rich seminal fluid provides strong contrast enhancement with Lugol’s iodine.

### Ductus deferens

NanoCT imaging shows the ductus deferens as paired muscular tubes with concentric hyperdense smooth muscle layers surrounding a narrow lumen (Fig 3C). The paired structures function to support sperm transport from the epididymis to the verumontanum. The distal ampullary glands appear as hyperdense alveolar structures at the base of the ductus deferens.

### Rhabdosphincter

The rhabdosphincter, or external urethral sphincter, is clearly visualized on contrast-enhanced nanoCT due to high Lugol’s iodine uptake within the circumferential striated skeletal muscle fibers (Fig 3D). Emerging caudal to the bladder neck and encasing the prostatic urethra, nanoCT may be used to assess sphincter integrity, thickness, and structure which may have implications for urinary continence.

### Prostate lobes

The murine prostate consists of four paired lobes anterior, ventral, dorsal, and lateral. Each are distinguishable on nanoCT imaging by their unique ductal morphology and contrast enhancement patterns. Lugol’s iodine staining highlights both glandular and stromal components. The anterior prostate displays uniformly hyperdense regions, attributed to abundant prostatic secretions and dense stromal smooth muscle, both of which enhance iodine uptake (Fig 4A). In contrast, the ventral prostate features hypodense ducts with large lumens and minimal intraglandular folding, reflecting sparse secretions and reduced stromal density (Fig 4B). The dorsal prostate is visualized as small, hypodense acinar clusters extending caudal the prostatic urethra into the membranous urethra and dorsally surrounding the inferior aspects of the seminal vesicles (Fig 4C). Finally, the lateral prostate, positioned between the ventral and dorsal lobes, comprises intermediate-sized ducts with moderate luminal folding and high contrast uptake (Fig 4D).

**Fig 4.**
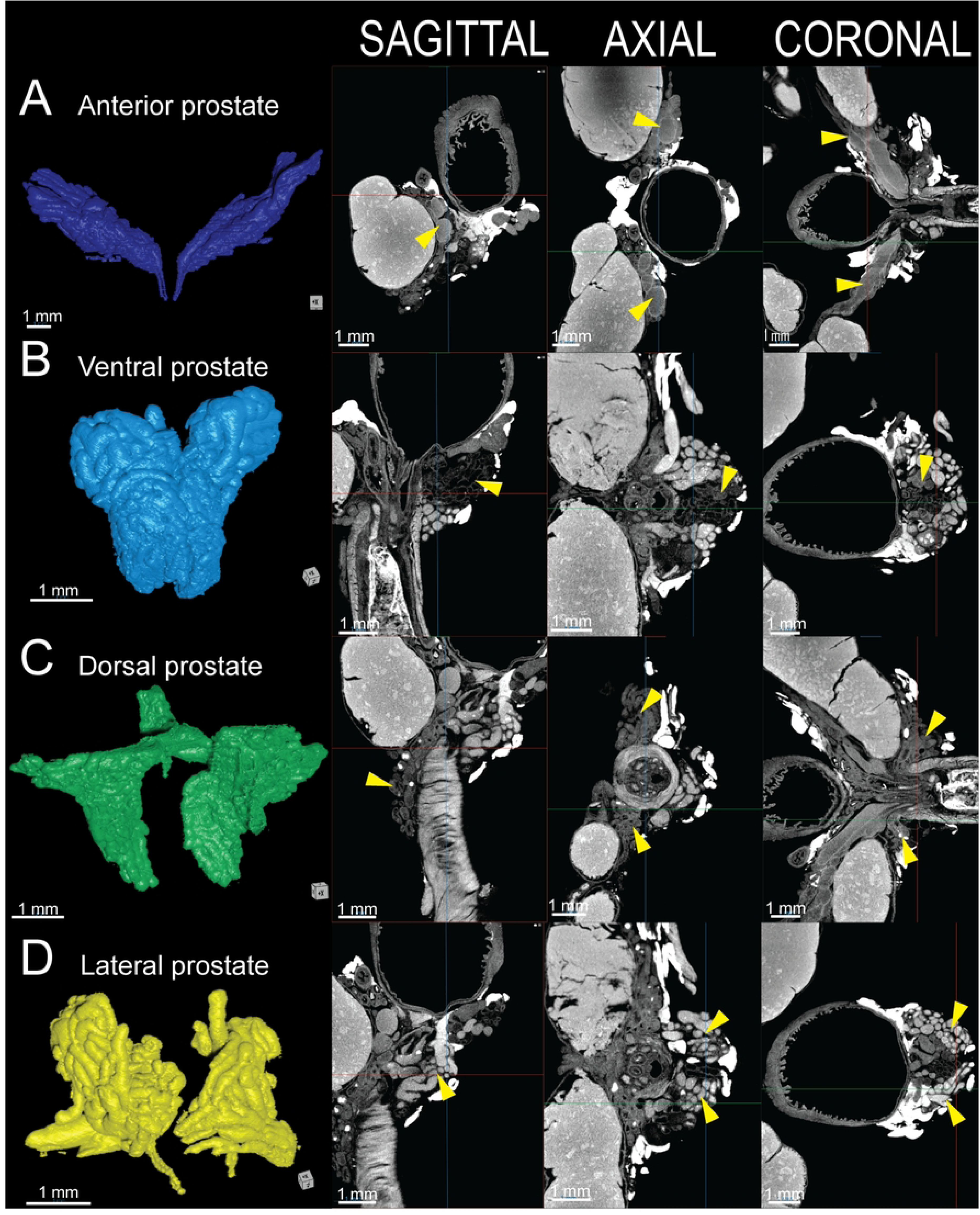
3D reconstruction and 2D tomographic views of the healthy 8-week-old C57BL/6J male murine prostate lobes. A) Anterior prostate (dark blue). B) Ventral prostate (light blue). C) Dorsal prostate (green). D) Lateral prostate (yellow). Next to each 3D reconstruction, corresponding sagittal (left), axial (center), and coronal (right) nanoCT slices depict the anatomical organization.

### NanoCT of the prostatic urethra

NanoCT imaging at the 3.5 μm/voxel spatial resolution provided visualization of the finer structures in the healthy 8-week-old C57BL/6J male murine prostatic urethra region. At this high spatial resolution, microscopic ductal architecture of the urethral lumen, seminal vesicle, ductus deferens, and prostate ducts as well as their branching patterns were able to be segmented, reconstructed, and observed (Fig 5A). In the region of the prostatic urethra, the urethral lumen begins as a narrow, elliptical structure just caudal to the bladder neck, gradually expanding distally as it receives secretions from the prostate and merges with the prostatic ducts (Fig 5B). NanoCT imaging revealed that the seminal vesicles contribute two ducts to the prostatic urethra (Fig 5C). Similarly, the paired ductus deferens contribute two ducts to the prostatic urethra (Fig 5D). These paired ducts maintain their spatial orientation along the length of the urethra, merging cranially to the verumontanum. Unlike in humans, where the seminal vesicle and vas deferens unite to form the ejaculatory ducts within the prostate, nanoCT imaging revealed that the murine ducts remain distinct until they converge at the verumontanum. High-resolution nanoCT imaging was able to visualize the individual ducts originating from the different prostate lobes. The anterior prostate lobe contributes two large, paired ducts (Fig 5E), which traverse medially from their lateral origins to merge with the superior aspect of the urethral lumen. These ducts do not undergo major branching as they extend cranially to caudally. In contrast, the ventral prostate lobe gives rise to 3–5 smaller ducts that enter the prostatic urethra (Fig 5F). These ducts, also oriented along the cranial-caudal axis, originate from the inferior-most aspect of the prostatic urethra, exhibiting some minor branching before merging with the urethral lumen. The dorsal and lateral prostate lobes contribute numerous ducts that enter the prostatic urethra (Fig 5G). These ducts are difficult to trace back to their lobe of origin due to extensive branching and tortuosity. As such, the ducts from both lobes are collectively referred to as the dorsolateral prostate ducts. These ducts exhibit complex expansion and distribution patterns before joining the urethral lumen at various levels.

**Fig 5.**
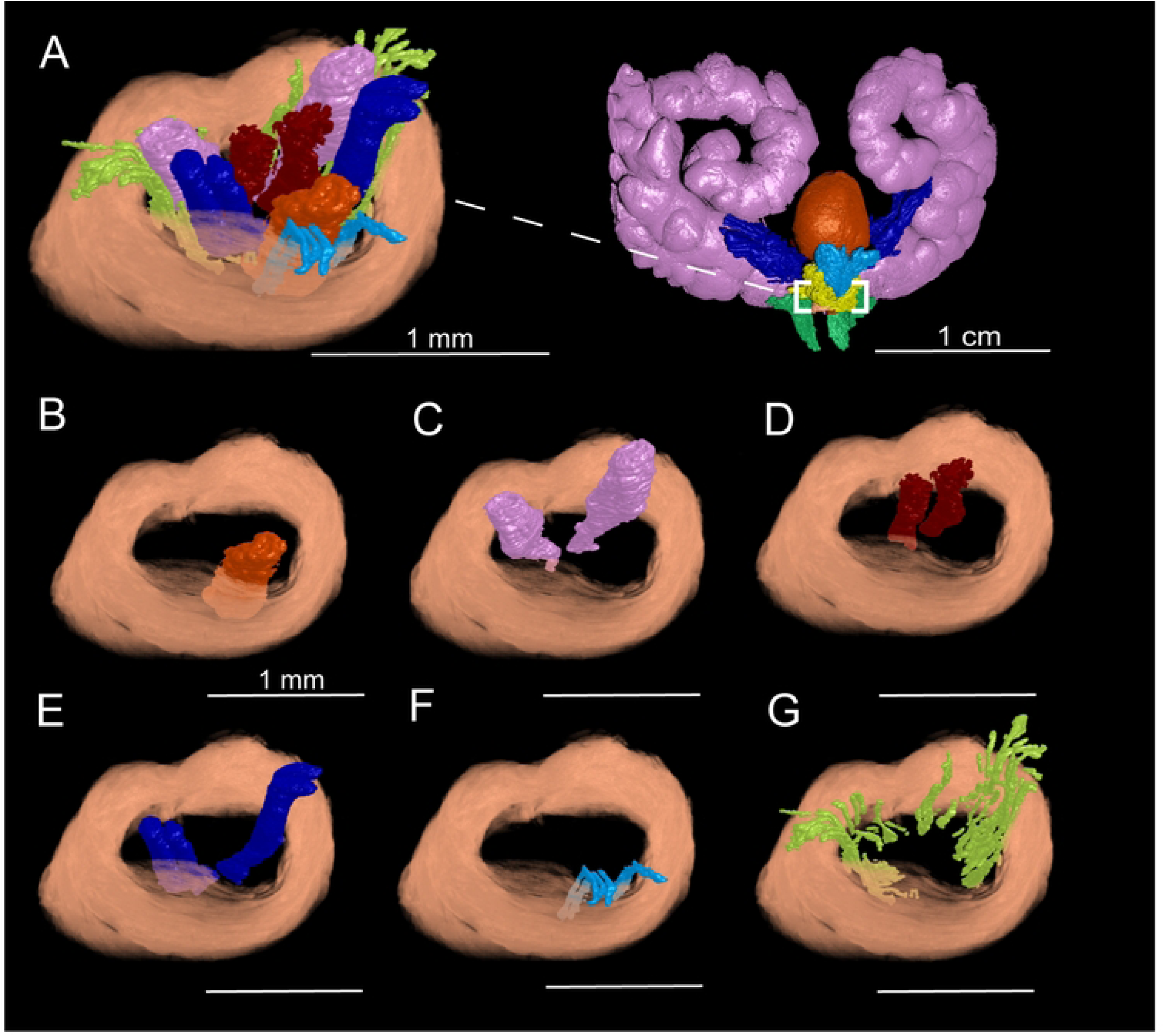
3D reconstruction of the healthy 8-week-old C57BL/6J male murine prostatic urethra. A) Isolated prostatic urethra (dashed box). B) Urethral lumen (orange). C) Seminal vesicle ducts (pink) D) Ductus deferens (red). E) Anterior prostate ducts (dark blue). F) Ventral prostate ducts (light blue). G) Dorsolateral prostate ducts (lime green).

The shape and structure of the prostatic urethra varies significantly along its length, influenced by the periurethral ductal system at each point. High-resolution nanoCT imaging allowed for visualization of this internal structure. 2D tomographs revealed that the rhabdosphincter begins approximately 200–300 μm distal to the bladder neck at the ventral aspect of the urethral lumen (Fig 6B). From this point, the muscle extends obliquely from ventral to dorsal, fully encircling the prostatic urethra (Fig 6D). Notably, this region (approximately 500-700 μm in length) marks the entry point for all primary ducts into the prostatic urethra: the urethral lumen (from the bladder), paired seminal vesicles, paired ductus deferens, and primary ducts from each prostate lobe. The point at which the rhabdosphincter fully encircled the prostatic urethra was the used here to define the anatomical start of the prostatic urethra region. Axial scans showed evidence of convergence of the paired anterior and ventral prostate ducts with the urethral lumen as early as 100-150 μm caudal to this landmark (Fig 6E and Fig 6F). To ensure consistency in our quantitative analyses, all measurements were performed 50 μm caudal to the point of rhabdosphincter closure. This location was selected to establish a reproducible anatomical landmark that captures the onset of the prostatic urethra while minimizing variability associated with more distal regions where the risk of omitting anatomical structures is higher.

**Fig 6.**
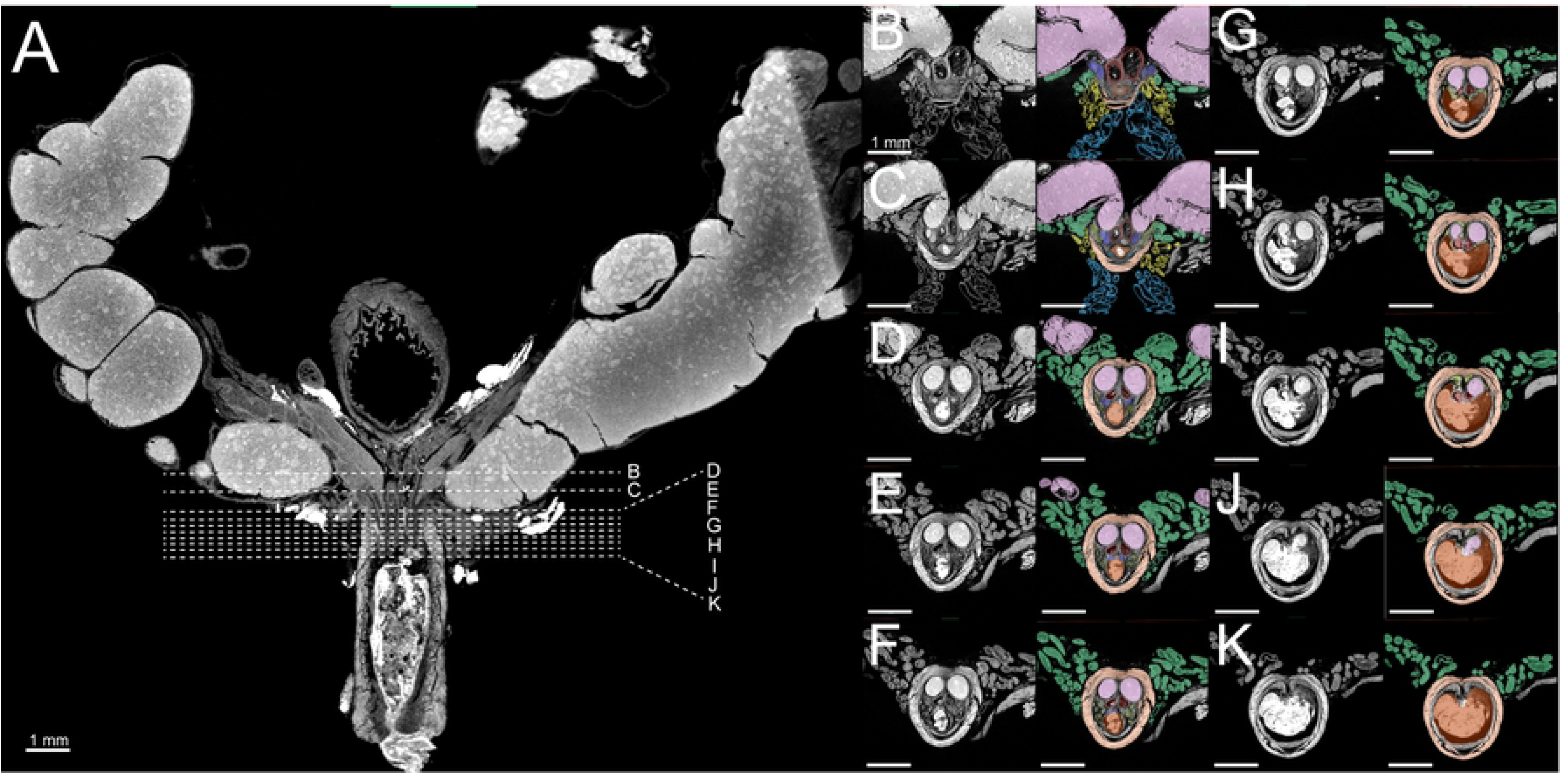
Axial Images of the of the healthy 8-week-old C57BL/6J male murine prostatic urethra. A) Coronal reference image indicating the approximate locations of axial slices. B-K) Serial axial nanoCT images of the prostatic urethra and surrounding structures along the cranial-caudal axis. Sections begin caudal to the bladder neck and extend to the verumontanum. Urethral lumen (orange), seminal vesicle (pink), rhabdosphincter (light orange), anterior prostate (dark blue), ventral prostate (light blue), lateral prostate (yellow), dorsal prostate (green), dorsolateral prostate ducts (lime green).

### Quantitative analysis of key features of the prostatic urethra

Contrast-enhanced nanoCT imaging of the murine prostatic urethra enabled for detailed measurements of key structural features *ex vivo* without the need for histologic sectioning. These key measurements are of particular interest in the study of lower urinary tract dysfunction. In this study, urethral lumen area, diameter, and perimeter, rhabdosphincter thickness, prostatic duct count, and prostatic urethral length were reported. A summary of prostatic urethra measurements are shown in Table 1. Urethral lumen area ranged from 0.07 mm^2^ to 0.16 mm^2^. Luminal diameter ranged from 0.21 mm to 0.39 mm. Luminal perimeter ranged from 0.79 mm to 1.44 mm. Rhabdosphincter thickness ranged from 0.1887 to 0.2375 mm. Prostatic duct number ranged from 28 to 47 ducts. Finally, prostatic urethra length ranged from 1.09 mm to 1.69 mm.

**Table 1.** Quantitative analysis of key features of the mouse prostatic urethra.

| Mouse | Urethral Lumen |  |  | Rhabdosphincter<br>thickness (mm) | Prostatic duct<br>count | Prostatic urethra length<br>(mm) |
| --- | --- | --- | --- | --- | --- | --- |
| | Area<br>($\text{mm}^2$) | Diameter<br>(mm) | Perimeter<br>(mm) | | | |
| 1 | 0.16 | 0.39 | 1.44 | 0.1887 | 32 | 1.69 |
| 2 | 0.07 | 0.35 | 0.99 | 0.2375 | 28 | 1.13 |
| 3 | 0.03 | 0.21 | 0.79 | 0.1925 | 31 | 1.49 |
| 4 | 0.09 | 0.36 | 1.16 | 0.2375 | 47 | 1.09 |

## Discussion

In this study, we provide a comprehensive, high-resolution 3D anatomical characterization of the murine male lower urogenital tract using nanoCT imaging. By preserving the spatial organization of key structures, particularly the prostatic urethra and associated prostate lobes, we present an anatomical framework that offers new insights into the mouse model’s relevance for studying human prostate biology and pathology, including lower urinary tract dysfunction (LUTD) and benign prostatic hyperplasia (BPH). This detailed anatomical mapping provides a foundation for further work into the role of prostatic architecture and metrics in disease development.

Our findings corroborate existing knowledge of the murine lower urogenital tract structure while adding a level of 3D detail that was previously unattainable with traditional methods such as histology.[27] NanoCT imaging enhanced visualization of the urinary bladder, seminal vesicles, ductus deferens, and prostate lobes, with a particular focus on the prostatic urethra. The ability to visualize ductal structures and their precise relationships to the urethral lumen is key to understanding pathologies like prostatic hyperplasia and LUTD. Unlike traditional high-resolution tissue reconstruction methods, which require embedding, sectioning, H&E staining, imaging, and slide by slide segmentation and reconstruction, nanoCT allowed us to reconstruct our tissues while maintaining native anatomical architecture in a significantly shorter time.[19, 20]

In both mice and humans, the periurethral zone posits itself an area of significant homology. In the human prostate, the transition zone, which encircles the proximal urethra and accounts for 5-10% of total prostate volume, is the primary site of BPH-related pathology.[28, 29] Anatomically, the human prostate transition zone consists of two regions whose ducts emerge from the posterolateral urethral wall, just proximal to the urethral orifice and the preprostatic sphincter, a collection of smooth muscle surrounding the proximal urethra.[30–32] These ducts extend laterally, curve anteriorly, and branch toward the bladder neck, with medial ducts penetrating the sphincter. Our nanoCT reconstructions reveal that the murine prostatic urethra region is surrounded by a dense network of ducts, primarily arising from the dorsal and lateral prostate lobes and to a lesser extent the anterior and ventral prostate. These ducts exhibit significant branching and direct connections to the urethral lumen, paralleling the human transition zone’s ductal architecture. Furthermore, like the fibromuscular capsule encasing the human prostate, the murine periurethral region is enclosed by thick circumferential muscle fibers comprising the rhabdosphincter. As periurethral ducts in the mouse prostatic urethra expand in size or number, due to aging or hormonal related changes, the rhabdosphincter may act as a mechanical barrier, restricting outward growth and directing expansion into the urethral lumen, thereby impinging on the urethral space.[9, 11] This mirrors the glandular and stromal hyperplasia in BPH that expands upon the urethral lumen, contributing to obstruction and LUTS.[30] These shared anatomical features, particularly ductal architecture, proximity to the urethral lumen, and periurethral organization, support the conclusion that the murine prostatic urethra closely parallels the human transition zone and should serve as the primary focus for studies of prostate-mediated lower urinary tract dysfunction in murine models. Our data further revealed that the anterior and ventral prostate lobes exhibit comparatively less communication with the urethral lumen and simpler ductal branching patterns. These lobes are anatomically positioned more distally from the urethral lumen, suggesting that they may play a less prominent role in modeling periurethral disease processes.

The ability for quantitative analysis of the prostatic urethra further strengthens the utility of nanoCT in future anatomical and pathological studies. We have previously shown that an MRI-based methodology can be used to quantify pathological changes within the murine lower urogenital tract.[11, 24, 25] However, this was limited to a resolution of 100 × 100 × 100 μm^3^, whereas our improved method achieves a spatial resolution of 3.5 × 3.5 × 3.5 μm^3^. Our data provides valuable baseline measurements of key structural features, including urethral lumen area, rhabdosphincter thickness, duct number, and prostatic urethra length. These measurements can serve as critical metrics for evaluating disease progression and the effects of therapeutic interventions in future studies of BPH and LUTD. Previous studies in the mouse have shown that an increase in prostatic duct number within the periurethral region is associated with lower urinary tract dysfunction.[9, 11] Therefore, duct count and branching morphogenesis can be studied in greater detail in the context of aging or treatment with antiproliferative agents (e.g. 5-α reductase inhibitors) and may lead to insights into how changes in in these structures contribute to urinary function.

In this study, we aimed to provide a high-resolution anatomical framework for studying the murine lower urogenital tract. The detailed 3D reconstructions and quantitative measurements presented here offer improved methods for exploring structural changes associated with BPH and LUTD in murine models. NanoCT is particularly valuable because it preserves native tissue architecture while providing higher spatial resolution than microCT or MRI, making it suitable for detailed anatomical imaging. However, several limitations remain. NanoCT is inherently an *ex vivo* imaging modality, which restricts its ability to assess longitudinal anatomical changes over the course of disease progression. In contrast, other imaging technologies like MRI, ultrasound, and optical methods enable *in vivo* imaging and can be used to monitor structural and functional changes in living animals over time.[33] For example, MRI offers adequate soft tissue contrast though it is limited by longer acquisition and image processing times. Ultrasound provides real-time visualization of superficial structures with reasonable spatial resolution but is limited by its inability to penetrate bone or air-filled tissues. Optical modalities such as bioluminescence and fluorescence imaging allow for high-throughput, transgene-based tracking of cell function or viability, though their utility is limited by low anatomical resolution and requires genetic manipulation of the animal model.[34] Moreover, while contrast enhancement is necessary for nanoCT imaging, it carries the potential to introduce artifacts or alter tissue integrity, although the use of Lugol’s iodine as an *ex vivo* contrast agent has not been shown to compromise downstream standard histological and immunohistochemical analyses.[35, 36] Finally, this study uses tissues from healthy, 8-week-old male C57BL6/J mice and does not consider anatomical variability due to age, sex, mouse strain, or pathology. Future studies will address these limitations and leverage this anatomical framework to investigate how such variables influence the murine lower urogenital tract. Overall, despite its limitations, nanoCT remains a powerful tool for high-resolution, high-throughput anatomical analysis relevant to prostate biology and disease research.

## Acknowledgements

We thank Dr. Gary Scheiffele and Dr. Edward Stanley from the University of Florida Herbert Wertheim College of Engineering Nanoscale Research Facility, and Dr. Karl Jepsen, Dr. Robert Goulet, and Andrea Clark from the University of Michigan NanoCT Core for their expertise, guidance, and overall assistance in scan optimization and image preprocessing. We would like to thank Sophia M. Vrba, Han Zhang, Avan N. Colah, and Alexis E. Adrian for their review of the manuscript.

## Notes

### Competing Interest Statement

The authors have declared no competing interest.

